# An *in vivo* model for transcranial direct current stimulation of the motor cortex in awake mice

**DOI:** 10.1101/2025.04.30.651337

**Authors:** Carlos Andrés Sánchez-León, Pablo Alejandro Reyes-Velasquez, Christoph van Thriel, Michael A. Nitsche

## Abstract

Transcranial direct current stimulation (tDCS) is a non-invasive brain stimulation technique mainly used in humans, in which weak direct currents are applied over the scalp to alter cortical excitability and induce neuroplasticity. Previous studies have demonstrated the value of tDCS for modulating sensory, motor, and cognitive functions, nevertheless, knowledge about how externally applied electric fields affect different components of neuronal networks is still incomplete, and *in vivo* animal models, which are required for a deeper understanding, are not fully developed. To evaluate the impact of tDCS on cortical excitability, many human experiments assess motor evoked potentials elicited by motor cortex (M1) stimulation. To develop a related *in vivo* animal model, we recorded electrical activity in M1 of alert mice during and after administration of tDCS over M1. M1 excitability was chronically recorded from layers 2-3, layer 5 and layer 6, evoked by stimulation of the ventral lateral nucleus of the thalamus (VAL). M1-tDCS was applied at 100 and 200 µA for 5 s to test the acute effects on neuronal excitability, and for 15 min to induce after-effects.

Acute M1-tDCS increased and decreased the amplitude of VAL-evoked potentials in a polarity-, layer- and intensity-dependent manner. For 15 minutes of anodal or cathodal tDCS, a similar polarity- and intensity-dependent modulation of VAL-evoked potential amplitudes during the 15 minutes of stimulation was observed. After tDCS was switched off, the highest intensity of anodal stimulation induced a significant excitability enhancement during at least two hours after stimulation, whereas the after-effects of cathodal tDCS were less pronounced.

The current study demonstrates the feasibility of a mouse model of M1-tDCS to accomplish similar modulatory effects of tDCS on cortical excitability as observed in human experiments. A proper adjustment of tDCS parameters, as compared to application in humans, is however required to obtain these translational effects.

## Introduction

Pre-clinical animal models are essential in translational neuroscience to investigate neurophysiological mechanisms of normal and pathological functioning of the human brain, and interventions targeting alterations of brain functions. By using this combination, Casaretto et al. (Casarotto et al., 2021) were able to identify the TRKB neurotrophin receptor as a long-time overlooked target of antidepressant (AD) drugs. Thus, manipulating central nervous system (CNS) functions by whatever means should be well understood for optimal and informed application in humans to make use of its full potential. Besides relevant methodological advances in the last decades to explore physiological foundations of psychological and behavioral processes in humans, the exploration of respective mechanisms at the cellular and molecular levels still faces important limitations. This applies also for revealing mechanisms of non-invasive brain stimulation (NIBS), which raised increasing interest in the last years for exploration of the physiological foundation of psychological and behavioral processes in humans, and consequently well-suited animal models reflecting human applications in this field are needed.

Transcranial direct current stimulation (tDCS) is a NIBS technique characterized by the application of constant electric currents over the scalp (Nitsche and Paulus, 2000) that are too weak to directly induce action potentials, but alter spontaneous neuronal activity, and excitability (Polanía et al., 2018; Stagg et al., 2018). Previous *in vivo* animal experiments demonstrated the capacity of externally applied currents to modulate spontaneous cortical activity and excitability (Creutzfeld et al., 1962; Bindman et al., 1964; Purpura and McMurtry, 1964; Cambiaghi et al., 2010; Sánchez-León et al., 2021, 2025). Moreover, in the last two decades, tDCS has demonstrated its feasibility to modulate neuronal excitability, and spontaneous activity during online stimulation, i.e., during the administration of external electric currents, most likely via an alteration of neuronal resting membrane potentials, but also to modulate cortical excitability, and induce synaptic plasticity after tDCS cessation also in humans (Nitsche and Paulus, 2001; Nitsche et al., 2005, 2008). The latter effects share features with long-term potentiation (LTP) and depression (LTD), such as calcium- and glutamatergic NMDA receptor dependency (Nitsche et al., 2003, 2004). For this reason, an increasing interest to investigate how tDCS modulates cognitive, and behavioral processes, as well as clinical symptoms in neurological, and psychiatric diseases has emerged (Brunoni et al., 2016; Woods et al., 2016; Bikson et al., 2019).

One of the most frequently used pre-clinical paradigms to study tDCS effects in humans is stimulation of the primary motor cortex (M1), where several studies have demonstrated (Nitsche and Paulus, 2000; Bikson et al., 2019) that when the anode is placed over M1 (anodal stimulation), tDCS enhances spontaneous activity and excitability of the target area, but when the cathode is placed over the same region (cathodal stimulation), spontaneous activity and excitability are reduced. These effects depend not only on stimulation polarity, but also on the specific orientation of the induced electrical field in relation to neuronal/ axonal orientation (Nitsche and Paulus, 2000; Rogalewski et al., 2004), because an efficient stimulation of cerebral neurons via constant electrical fields depends on the alignment of the electrical field to the longitudinal axis of respective neurons (Kabakov et al., 2012; Rahman et al., 2013). Furthermore, depending on the duration (Nitsche and Paulus, 2001; Agboada et al., 2019; Mosayebi Samani et al., 2019) and intensity (Jamil et al., 2017; Agboada et al., 2019; Mosayebi Samani et al., 2019) of stimulation, the excitability alterations observed under a specific stimulation polarity (anodal or cathodal) can be potentiated, but also reversed, most likely because of the calcium dynamics presumed to underly neuroplastic effects, including those elicited by tDCS (Lisman, 2001; Nitsche et al., 2003). Understanding the neural basis underlying this dosage-, and protocol-dependency of stimulation effects in detail is essential to develop new strategies to optimize the effects of tDCS in humans.

Similar to other areas in cognitive and clinical neuroscience and psychopharmacology, pre-clinical animal models are a potentially powerful tool to fully understand neurophysiological changes, including their magnitude and duration during and after tDCS (Sánchez-León et al., 2018b, 2018a). *In vivo* animal models offer the opportunity to study the impact of tDCS on intact active brains, allowing the direct measure of intracranial neuronal excitability, and activity with high temporal, spatial and molecular resolution (Cambiaghi et al., 2011; Márquez-Ruiz et al., 2012; Fernández et al., 2021). Regarding tDCS, animal models have played a key role in elucidating some of the mechanisms, and brain features involved in its effects, such as neuronal morphology (Radman et al., 2009), orientation of the somatodendritic axis with respect to the electric field (Bikson et al., 2004; Sánchez-León et al., 2025), the effects on different cortical layers (Sun et al., 2020), or the involvement of astrocytic, and other glial cells (Monai et al., 2016; Mishima et al., 2019). Nonetheless, most of these studies were conducted under anesthesia (Cambiaghi et al., 2010; Monai et al., 2016; Liu et al., 2019), or in reductionistic slice preparations (Bikson et al., 2004; Rahman et al., 2013; Kronberg et al., 2020), which might interfere with tDCS-induced plasticity and complicate the interpretation of results. In this context, the establishment of an awake mouse model for non-invasive electrical stimulation over M1 is crucial to foster the translatability of mechanisms of action between human, and animal studies.

In this study, we show how to combine simultaneous *in vivo* electrophysiological recordings in awake mice from different cortical layers, and tDCS over M1. For that, we evoked M1 activity by electrical stimulation of the ventro-antero-lateral nucleus of the thalamus (VAL) while testing M1-tDCS protocols at different intensities (100 and 200 µA) and different intervention durations, including short stimulation (5s) to explore online effects, and longer stimulation (15 minutes) to investigate neuroplastic alterations. For online administration of tDCS, we hypothesized an intensity- and polarity-dependent modulation of evoked potentials, with increased excitability during anodal tDCS and decreased excitability during cathodal stimulation, and stronger effects with larger stimulation intensities. Regarding neuroplastic effects, we hypothesized identically directed polarity- and intensity-dependent effects.

## Methods

### Animals

Experiments were carried out on a total of 25 adult wild-type mice (C57BL/6JRj, Janvier Labs, France; 10-30 weeks old, both males and females), 15 for immediate effect experiments and 20 for after-effects, 10 of them overlapping between experiments. Details about the allocation of the mice and gender distribution are given in the results section. Experimental procedures were carried out in accordance with European Union guidelines (2010/63/CE) and approved according to German regulations (LANUV, Akz.: 81-02.04.2020.A208) for the use of laboratory animals.

### Surgery

Prior to the experiments, stereotaxic surgery was performed including craniotomies for intracortical electrode insertion, implantation of the tDCS electrode and the headplate for head-fixed experiments. Mice were deeply anesthetized with a ketamine–xylazine mixture (ketamine 100 mg/ml, Bayer Health Care, Leverkusen, Germany; xylazine, 20 mg/ml, Pfizer, Berlin, Germany) and kept on a heating pad to maintain constant body temperature at 37°C. In addition, buprenorphine (Bupresol 0.3 mg/ml, CP-Pharma, Burgdorf, Germany) was given perioperatively to provide post-operative analgesia. A midline incision was made to expose the skull and two marks were made over the bone centered over the right M1 (AP = +1 mm; L = -1 mm; relative to bregma) and right ventro-antero-lateral thalamus (VAL) (AP = -1.3 mm; L = -1 mm; relative to bregma). Three small screws were placed on the skull, two on either side of the cerebellum and a third over the left parietal bone, and a thin stainless steel headplate was placed just behind the lambda suture and fixed to the screws and skull using Superbond (Hentschel-Dental, Teningen, Germany). After that, a custom-made ring-electrode (see section electrode fabrication) was placed over the skull centered over the right M1 mark and fixed with a layer of Loctite super glue (Henkel, Düsseldorf, Germany) and then dental cement (DuraLay, Illinois, USA), making sure not to pour these substances between the electrode and the skull. Once fixed, two holes (1.5 mm ø) were drilled in the previously marked bone to allow access to M1 and the motor thalamus. In addition, a silver reference electrode (see section electrode fabrication) was implanted over the dura surface under the left frontal bone (AP = +1 mm; L = +2 mm; relative to bregma). Finally, the exposed dura mater surface was protected with fast curing silicone elastomer (Kwik Sil, WPI, Florida, USA) until recording sessions. Buprenorphine (0.05 mg/kg) was administered for the next three days every 8-12 hours to provide analgesia.

### Electrode fabrication

For the tDCS ring-electrode, a silver wire (bare silver wire, ø: 635 µm; A-M Systems, Washington, USA) was cut into pieces of 1 cm length and one end was curved and welded to form a closed loop. Subsequently, this end was pressed with pliers to flatten the surface and create a 2.5 mm inner ø, 4 mm outer ø stimulation surface that was subsequently chlorinated. To insulate the shaft of the electrode, it was introduced into a flexible tubing sleeve (inner ø: 0.508 mm; outer ø: 0.939 mm; wall thickness: 0.215 mm; Silicone Tubing, A-M Systems, Washington, USA). Finally, the non-curved end of the electrode was soldered to a connector pin.

For the recording reference electrode, a silver electrode (bare silver wire, ø: 381 µm, A-M Systems, Washington, USA) was cut into pieces of 1 cm length, a loop (2 mm ø) was made at one end to facilitate posterior grasping by the pre-amplifier system and the opposite end of the electrode was braided and filed to avoid damaging the dura mater.

### Electrical stimulation

To elicit evoked activity in M1, a brief electrical pulse was delivered to the VAL nucleus. For this, an implantable bipolar stimulation electrode (100 µm quartz-glass insulated platinum tungsten fiber, impedance below 150 kOhm, protrusion: 6 and 5.5 mm, Thomas Recording, Giessen, Germany) was lowered through the thalamic craniotomy to the level of the VAL nucleus. The electrical stimulus consisted of a single square pulse (250 µs; 0.05-0.5 mA), stimulation intensity was adjusted to obtain an evoked potential with half of the maximum amplitude and delivered by an isolation unit (ISU-175, Cibertec, Madrid, Spain) connected to a digital stimulator device (CS420, Cibertec, Madrid, Spain).

For tDCS administration, two different current intensities (100 and 200 µA) were applied between the ring-electrode over M1 and a surface electrode consisting of a rubber rectangle (6 cm^2^) attached to the back of the mice and moistened with electrode gel (electrogel, Electro-Cap International, Ohio, USA). To characterize the online effects induced by tDCS, during the recording session tDCS was applied repetitively for a short duration (15 s, including 5 s linear ramp-up and down at the start/end of stimulation, separated by 10 s without stimulation) at 2 randomly interleaved intensities (100 and 200 μA). Thalamic stimulation was delivered during the no-stimulation period 1s before tDCS began and during tDCS 1s before tDCS ramp-down began. 15 trials for each tDCS intensity and polarity were applied. To induce tDCS after-effects, after 30 min of baseline measures of M1 excitability, tDCS was delivered over M1 for 15 min (30 s linear ramp-up and down before and after the 15 min of stimulation) for anodal or cathodal stimulation, and for 90 s (30 s linear ramp-up followed by 30 s at 200 μA and 30 s linear ramp-down followed by 14.5 min without stimulation) for sham stimulation. Thalamic stimulation was delivered every 10 ± 2 s during the whole session for up to two hours after the end of tDCS. The different protocols for transcranial direct current application were designed in Spike2 (Cambridge Electronic Design, CED, Cambridge, UK), and sent to a battery-driven linear stimulus isolator (WPI A395, Florida, USA) through an analog output from the acquisition board (CED 1401-3A, Cambridge, UK).

### Intra-cortical recording

Mice were habituated to the head-restraint for two days and any recording session began at least 48 hours after the last buprenorphine administration to avoid interference of the drug with neuronal excitability, and stimulation effects. The animals were placed on a treadmill with a rotatory encoder (Janelia Experimental Technology, Virginia, USA) for locomotion monitoring and the head was fixed to the recording apparatus. Silicone elastomer was removed with the aid of a surgical microscope, and the exposed cortical surface was carefully cleaned with a super fine forceps (Dumont #5, FST, Heidelberg, Germany), and cotton swab without damaging the dura mater. The implantable recording electrode array (three consecutive shafts each 100 µm quartz-glass insulated platinum tungsten fiber electrodes, impedance 0.5 - 0.8 mOhm, protrusion: 2.2, 2.7 and 3mm, Thomas Recording, Giessen, Germany) was lowered (MMO-220A, Narishige, Tokyo, Japan) into the burr holes of the M1 craniotomy until the tips of the electrodes were around 0.2, 0.7 and 1 mm below the dura surface, and the reference wire electrode was attached to the recording reference electrode positioned at the contralateral hemisphere. The thalamic stimulation electrode was lowered (MMO-220A, Narishige, Tokyo, Japan) approximately 3.5 mm into the boreholes of the thalamic craniotomy surface, and the electrical stimulus was delivered every 10 ± 2 s to elicit a neuronal response in M1. The depth of both electrodes was adjusted until evoked potentials (EP) were clearly identified based on their waveform (occurrence of a reliable negative deflection) and latencies (with the first evoked neuronal activity appearing with a latency lower than 2 ms after the thalamic stimulus), and afterwards cemented together with the head holding system. All recording signals were amplified by a factor of 1.000 (PGMA-2 amplifier connected to a PA-16 pre-amplifier, Thomas Recording, Giessen, Germany) and digitized at 25 kHz (CED 1401-3A, Cambridge, UK). The remaining non-neuronal signals (tDCS and treadmill movement) were sampled at 5 kHz.

We then waited one week before we started experimental recordings to allow the electrodes and the tissue to stabilize. For the recordings, mice were placed on the treadmill and the head was fixed to the recording apparatus. We let the mice accommodate for 15 minutes to the head-fixed condition and then proceeded with the experimental protocols. One week was allowed between sessions to avoid interference of potential long-term plasticity effects in consecutive sessions, and the session order of for the after-effect experiments was randomized with respect to stimulation polarity and intensity.

### Data analysis

Data were analyzed in Spike2 (CED, Cambridge, UK) and Matlab (MathWorks Inc., Massachusetts, USA) using custom written software. SPSS (version 25, IBM, Armonk, NY)) and Matlab (MathWorks Inc., Massachusetts, USA) were used for statistical analysis. Evoked potentials (EPs) coincident with animal movement as well as electrical artifacts were removed from the analysis based on visual inspection. EP amplitudes were computed as the voltage difference between the first negative peak and the voltage value at initiation of the negative deflection. Latency was determined as the time interval from VAL stimulation to the main negative peak value.

For characterization of the evoked local field potentials in M1 in response to thalamic stimulation, event-related potential (ERP) analysis was performed in the EEGLAB toolbox. Data were segmented in epochs of 1 s using VAL stimulation as a trigger and baseline correction was done by subtracting the mean voltage level in the 500 ms preceding VAL stimulation.

For online effect experiments, we first calculated the mean amplitude of the EP per individual mouse for each factor combination (INTENSITY, POLARITY, STIMULATION and LAYER).

Then we conducted an ANOVA, with EP amplitude as dependent variable and INTENSITY (2 levels: 100 or 200 μA), POLARITY (2 levels: anodal or cathodal), STIMULATION (2 levels: tDCS ON or OFF) and LAYER (3 levels: 2-3, 5 and 6) as within-subject factors. After that, we normalized the EP amplitude during tDCS (tDCS ON) to baseline EP (tDCS OFF) 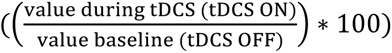 and applied another ANOVA with normalized EP amplitudes as dependent variable and INTENSITY, POLARITY and LAYER as within-subject factors, as described above. In case of significant effects of the ANOVAs, Fisher’s LSD post hoc tests were conducted.

For the after-effect experiments, EP amplitudes were averaged in 15 min time windows per individual mouse, to better define the time course of plasticity induction, and were normalized to the last baseline period (15 minutes before the initiation of tDCS). Subsequently, a mixed-model ANOVA was conducted, using normalized EP amplitudes as dependent variable, CONDITION (5 levels: anodal 100 and 200 μA, cathodal 100 and 200 μA and sham) as between-subject factor, since not all mice received all intervention conditions (sham (10 mice (6 females)), 100 μA anodal (11 mice (5 females)), 100 μA cathodal (14 mice (6 females)), 200 μA anodal (12 mice (6 females)), or 200 μA cathodal (12 mice (4 females))), and LAYER (3 levels: 2-3, 5 and 6) and TIME (11 levels: two periods before tDCS, one during tDCS and eight after tDCS) as within-subject factors. To compare tDCS effects within different polarities, we performed subsequent ANOVAs separately for cathodal and anodal stimulation, with INTENSITY (3 levels: sham,100 or 200 μA) of these stimulations as between-subject factor, and LAYER (3 levels: 2-3, 5 and 6) and TIME (11 levels) as within-subject factors. Post hoc comparisons were performed for every time point using Fisher’s LSD post hoc tests in case of significant results of the ANOVA tests. For all ANOVAs, Mauchly’s test of sphericity was conducted, and the Greenhouse-Geisser correction was applied when necessary. Statistical significance was set for all statistics at p < 0.05. For all analyses, the results are shown as mean ± SEM.

## Results

### Characterization of evoked potentials over M1 in response to thalamic stimulation

To obtain an electrophysiological biomarker of neuronal excitability in awake mice that can later be used to investigate and analyze the tDCS effects, we first recorded and characterized local field potentials in three different cortical layers of M1 (layer 2-3, layer 5 and layer 6) elicited by a brief electrical stimulation of a source of M1 afferences in the motor thalamus (i.e., VAL nucleus; Fig. 3A,B). The VAL stimulation (0.05-0.5 mA) evoked a short-latency neuronal response in M1 (event-related potential (ERP) from n = 10 mice; Fig. 3C) characterized by a main negative peak that appeared earlier in deeper layers (L2-3: 9.08 ± 0.61 ms; L5: 8.14 ± 0.58 ms; L6: 6.25 ± 1.01 ms), followed by a positive deflection in the two deeper layers. Next, we assessed the intensity dependency of the electrical stimuli applied to VAL and the amplitude of the main negative component. For that, we applied VAL stimulation at different intensities (0.05 to 0.5 mA at increasing steps of 0.05 mA) and normalized ERP amplitudes values by the amplitude achieved with maximum VAL stimulation intensity (0.5 mA; Fig. 3D). We observed a linear stimulation intensity-dependent increase of ERP amplitudes, as shown for an exemplary mouse in Fig. 3D and for averaged data of two mice in Fig 3E (L2-3: R^2^ = 0.93, p < 0.001; L5: R^2^ = 0.86, p < 0.001; L6: R^2^ = 0.85, p < 0.001.). This outcome allowed us to adjust the EP amplitude in each of the following experiments to elicit an EP with half of the maximum amplitude to guarantee the option to observe an increase or decrease of its components during and after tDCS intervention.

### tDCS online effects on M1 excitability

To explore tDCS effects during online stimulation of M1, we recorded EPs induced by VAL stimulation during anodal or cathodal tDCS in 15 mice (7 females). We applied tDCS pulses of 5 s duration (15 s in total, including 5 s linear ramp-up and down) at 2 randomly interleaved intensities (100 and 200 μA) preceded by a 10 s baseline period (Fig. 1). For any given intensity, the first pulse was anodal tDCS followed by 10 s of no stimulation and then a cathodal tDCS pulse at the same intensity. Thalamic stimulation was delivered during the baseline period 1s before tDCS began and during tDCS 1s before tDCS ramp-down began. At least 15 trials for each tDCS intensity and polarity were applied. For a representative animal (Fig. 4A), we show the averaged EP (n = 24 trials) in the baseline condition before tDCS began (dotted or dashed lines for EPs before anodal or cathodal tDCS, respectively), and superimposed are the recordings during anodal (red traces) or cathodal (blue traces) tDCS. First, we compared the values during baseline (tDCS OFF) with the values during tDCS (tDCS ON). The ANOVA analysis (Table 1) revealed significant main effects of polarity (d.f.: 1, F = 12.132, p = **0.004**) and layer (d.f.: 1.224, F = 5.784, p = **0.023**) but not of tDCS (d.f.: 1, F = 1.528, p = 0.237) or intensity (d.f.: 1, F = 4.150, p = 0.061).

**Figure 1.**
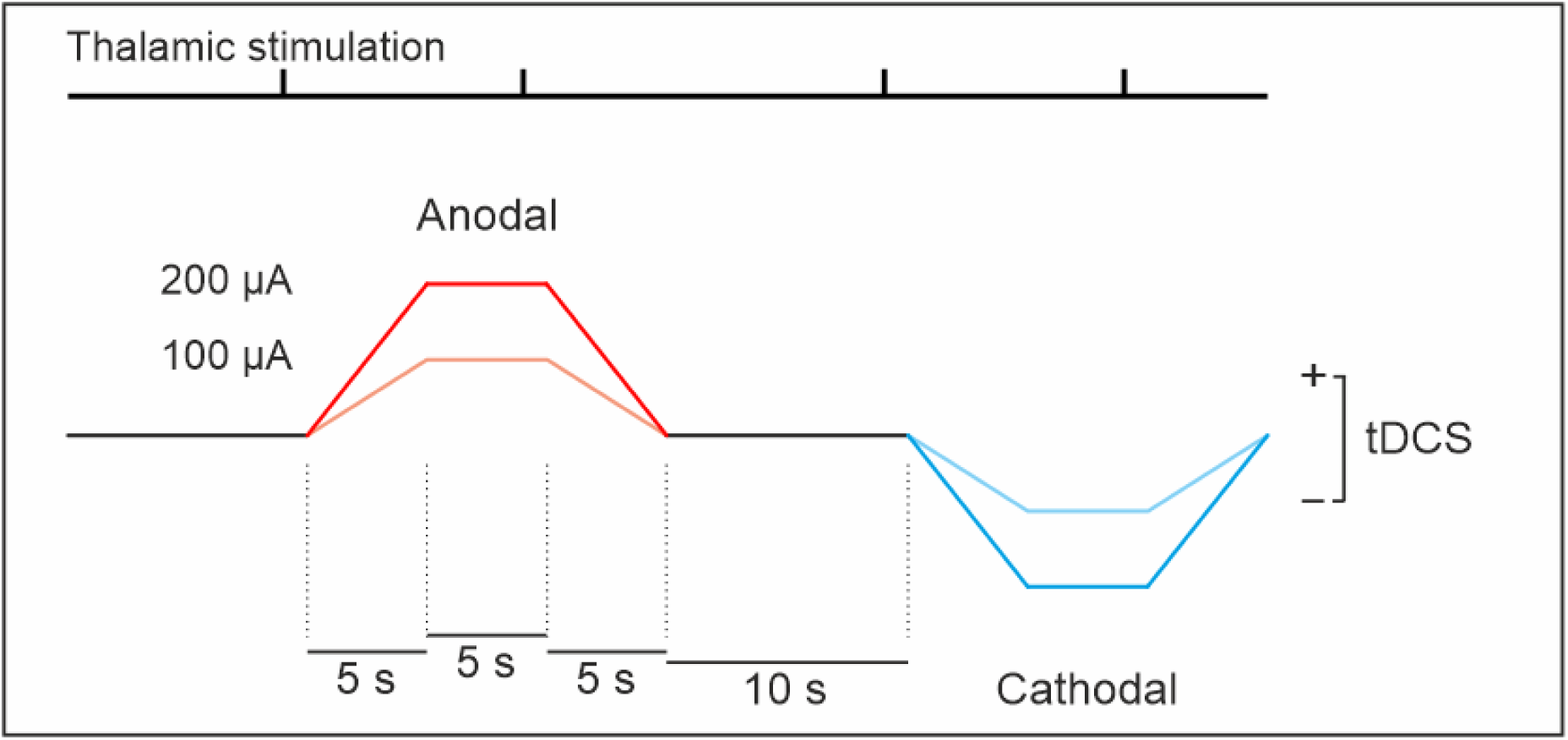
Experimental paradigm used to assess online effects. M1 evoked potentials were recorded before and during the administration of brief tDCS pulses of anodal (red lines) and cathodal (blue lines) stimulation. tDCS was applied for a short duration (15 s, including 5 s linear ramp-up and down at the start/end of stimulation, separated by 10 s without stimulation) at 2 randomly interleaved intensities (100 and 200 μA). Thalamic stimulation was delivered during the no-stimulation period 1s before tDCS began and during tDCS 1s before tDCS ramp-down began.

**Table 1.** Overall ANOVA conducted for the online effects of tDCS. The absolute evoked-potential amplitude values were compared in a 4-factorial repeated-measures ANOVA. d.f. = degrees of freedom

| Factor | d.f. | F-value | p-value | Partial Eta squared |
| --- | --- | --- | --- | --- |
| layer | 1.224* | 5.784 | **0.023** | 0.292 |
| polarity | 1 | 12.132 | **0.004** | 0.464 |
| stim | 1 | 1.528 | 0.237 | 0.098 |
| intensity | 1 | 4.150 | 0.061 | 0.229 |
| stim*polarity | 1 | 10.458 | **0.006** | 0.428 |
| stim*layer | 1.427* | 0.450 | 0.579 | 0.031 |
| stim*intensity | 1 | 0.714 | 0.412 | 0.049 |
| layer*polarity | 1.131* | 4.088 | 0.056 | 0.226 |
| layer*intensity | 1.263* | 4.521 | **0.040** | 0.244 |
| polarity*intensity | 1 | 6.501 | **0.023** | 0.317 |
| layer*polarity*stim | 1.126* | 4.536 | **0.046** | 0.245 |
| intensity*polarity*stim | 1 | 7.702 | **0.015** | 0.355 |
| layer*polarity*intensity | 1.297* | 5.314 | **0.026** | 0.275 |
| layer*stim*intensity | 1.469* | 1.388 | 0.266 | 0.090 |
| layer*polarity*stim*intensity | 1.082* | 6.698 | 0.019 | 0.324 |

Notes: \*: Greenhouse-Geisser corrected dfs

In accordance with the observation that anodal and cathodal polarities modulated EP amplitudes in opposite directions, the stimulation*polarity interaction (d.f.: 1, F = 10.458, p = **0.006**) showed a significant effect of stimulation whose direction depended on the polarity applied. Furthermore, we found an intensity-dependent effect, as shown by the significant interactions between layer*intensity (d.f.: 1.263, F = 4.521, p = **0.040**) and polarity*intensity (d.f.: 1, F = 6.501, p = **0.023**). Lastly, we also observed an intensity- and layer-dependent modulation of the polarity-specific effects on the EP amplitude (interaction layer*polarity*stim: d.f.: 1.126, F = 4.536, p = **0.046**; interaction intensity*polarity*stim: d.f.: 1, F = 7.702, p = **0.015**; interaction layer*polarity*intensity: d.f.: 1.297, F = 5.314, p = **0.026**; interaction layer*polarity*stim*intensity: d.f.: 1.082, F = 6.698, p = **0.019**). Post-hoc comparisons (Fig. 4B) between EP amplitudes during tDCS (tDCS ON) and baseline values (tDCS OFF) for all layers, intensities and polarities showed that anodal tDCS significantly increased the EP amplitudes for all intensities and layers, and that cathodal tDCS generated a significant decrease of EP amplitudes for all intensities for all layers (N = 15 mice). In addition, an intensity-dependent effect only during tDCS (tDCS ON) for layers 2-3 and 5 and for anodal and cathodal stimulation, but not for layer 6 ermerged.

As shown in Table 1, there was also a significant main effect for the factor layer (d.f.: 1.224, F = 5.784, p = **0.023**), and in Fig. 4A and B smaller EP amplitudes in deeper layers also in baseline conditions (tDCS OFF) were observed. For this reason, we conducted an additional ANOVA with normalized amplitude values 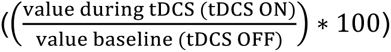 (Fig. 5 and Table 2) for each layer, polarity and intensity. As expected, a significant main effect for polarity (d.f.: 1, F = 70.086, p = <**0.001**) emerged, as well as a significant layer*polarity interaction (d.f.: 2, F = 5.802, p = **0.008**), demonstrating that the tDCS-induced modulation of EP amplitudes across layers is not uniform. In addition, we also observed different effects of tDCS depending on stimulation intensity, and layer (polarity*intensity: d.f.: 1, F = 25.318, p = <**0.001**; layer*polarity*intensity: d.f.: 2, F = 4.912, p = **0.015**). Post-hoc comparisons between the different layers (Fig. 5A, top for anodal, bottom for cathodal) showed a weaker modulation for the deeper layer 6 in comparison with layer 2-3 and 5 for anodal stimulation at 200 µA and layer 5 for cathodal tDCS at 100 and 200 µA. In addition, comparisons between the different tDCS intensities (Fig. 5B, top for anodal, bottom for cathodal) showed stronger effects for the higher tDCS intensity, resulting in increased EP amplitudes for anodal, and decreased MEP amplitudes for cathodal tDCS for 200, as compared to 100 µA in layers 2-3 and 5, but not in layer 6.

**Table 2.**
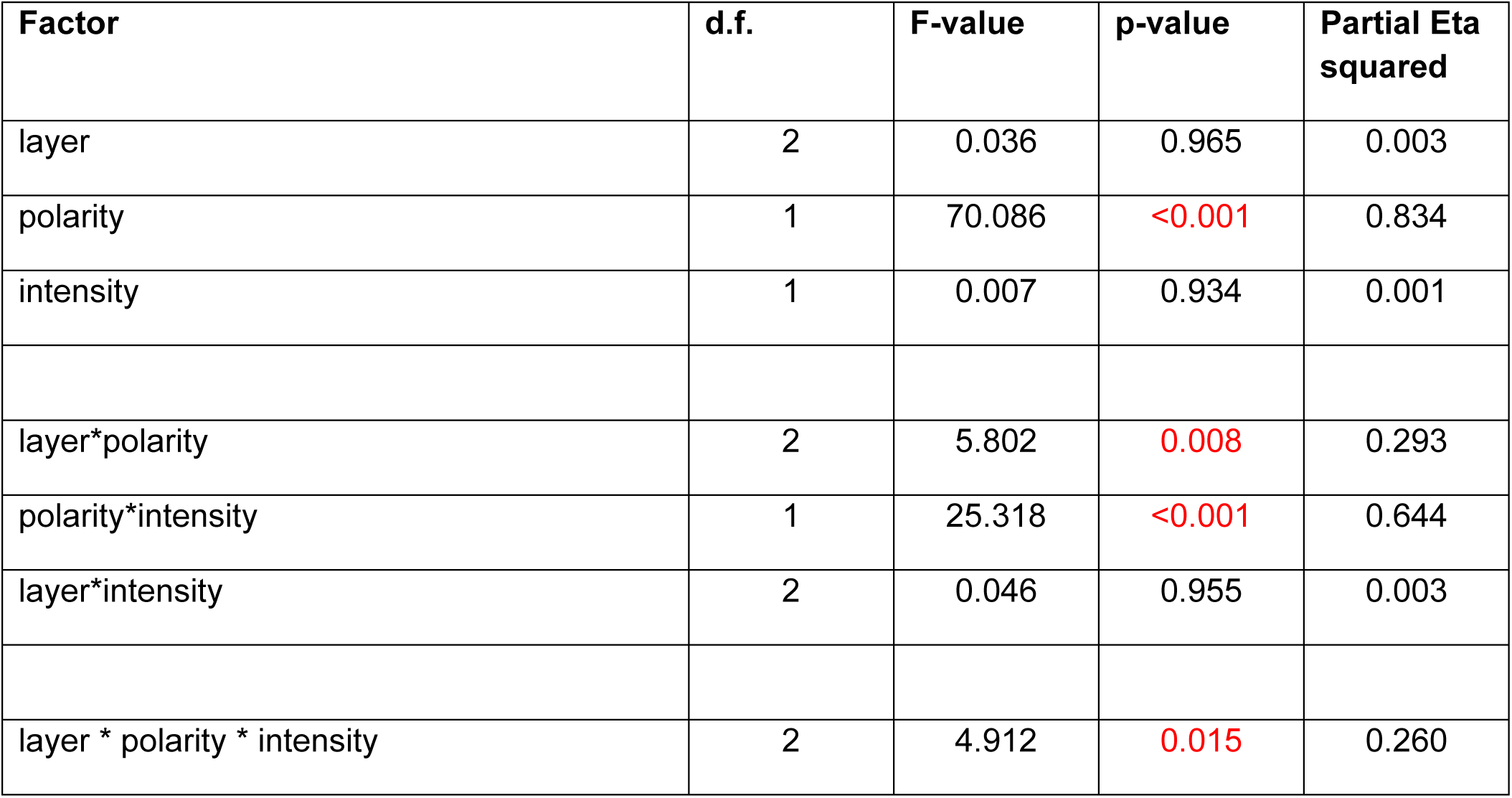
ANOVA conducted for the online effects of tDCS with normalized EPs. Normalized evoked-potential amplitude values were compared in a 3-factorial repeated-measures ANOVA. d.f. = degrees of freedom.

These results demonstrate the efficacy of M1-tDCS for online modulation of neuronal excitability, showing polarity-, intensity-, and layer-dependent modulation, with anodal tDCS increasing, and cathodal tDCS decreasing EP amplitudes and higher intensities eliciting stronger modulation for layers 2-3 and 5. Next, we explored if prolonged stimulation induced long lasting effects on M1 excitability that exceed the stimulation period.

### tDCS after-effects on M1 excitability

For these experiments, EP were assessed for 30 minutes for baseline recordings, during the following 15 minutes of tDCS at different intensities (sham (10 mice (6 females)), 100 anodal μA (11 mice (5 females)), 100 μA cathodal (14 mice (6 females)), 200 μA anodal (12 mice (6 females)), or 200 μA cathodal (12 mice (4 females))) and finally for 120 minutes after tDCS to check for after-effects. VAL stimulation was applied every 10 ± 2 seconds to avoid any anticipatory activity and/or plasticity due to regular repetition intervals (Fig.2). For each recording, EP were averaged in 15 min time windows. EP amplitudes were stable in the sham condition in all layers during the 165 minutes of the experiment (Fig.6, black circles), although we noticed a trendwise reduction of the EP amplitude in layer 2-3 (zoom in Fig.6A, sham, black trace) and increase in layer 6 (zoom in Fig.6C, sham, black trace). We first compared the normalized EP amplitude values by an overall ANOVA (Table 3). The results did not show a significant main effect of the factor layer, but significant main effects of Time (d.f: 2.754, F = 4.576, p = **0.005**), and Condition (d.f: 4, F = 5.001, p = **0.002**), and significant Time*Condition (d.f.: 11.017, F = 15.532, p = <**0.001**) and Layer*Time*Condition interactions were observed (d.f.: 18.073, F = 1.932, p = **0.014**).

**Figure 2.**
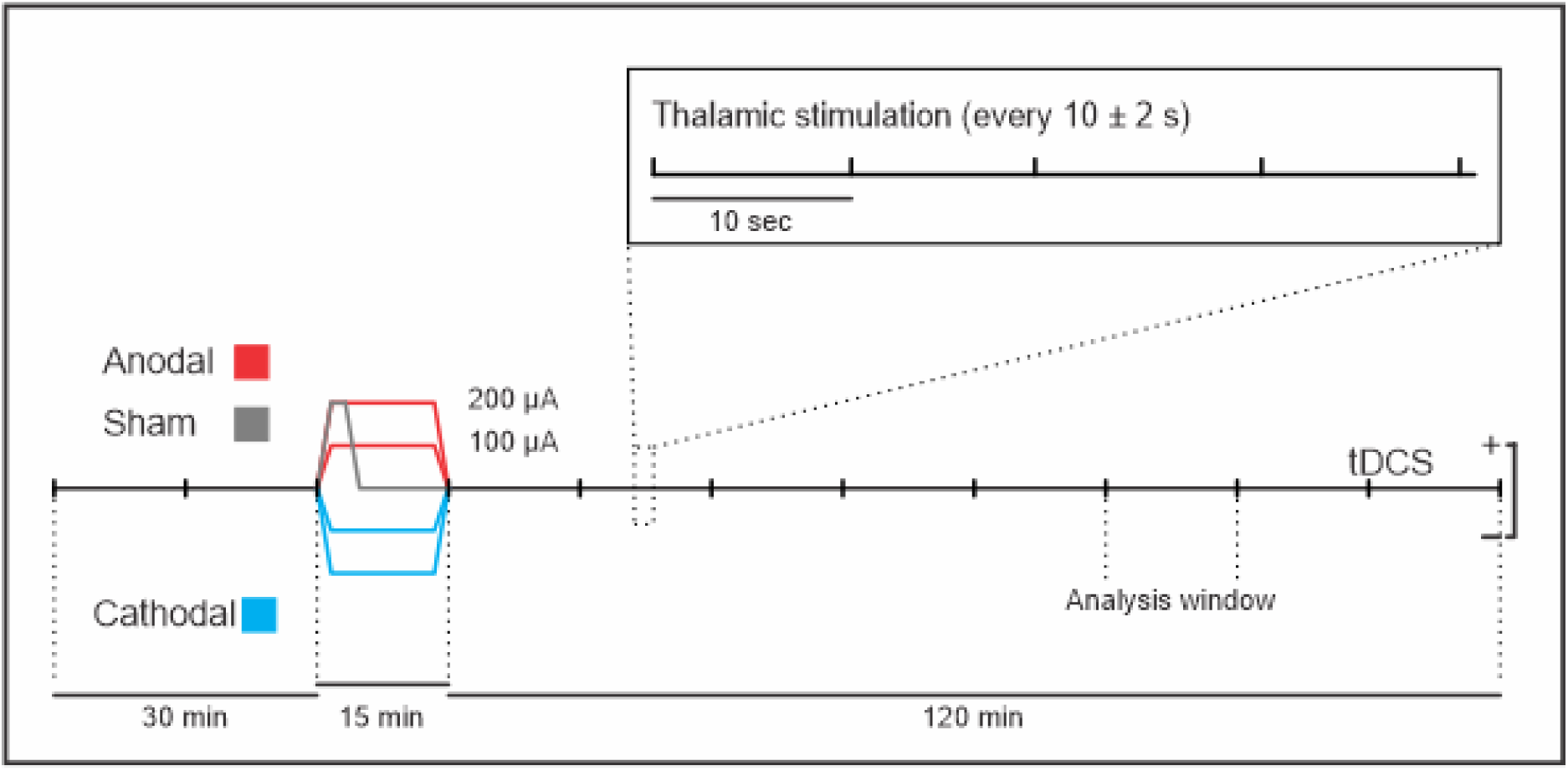
Experimental paradigm used to assess after-effects of DC stimulation. M1 Evoked potentials were recorded every 10 ± 2 s during the whole session. The protocol consisted of 30 min baseline EP measures followed by 15 min (30 s linear ramp-up and down before and after the 15 min of stimulation) of anodal (red lines) or cathodal (blue lines) tDCS at different intensities (100 and 200 μA) or sham stimulation (grey line) for only 90 s (30 s linear ramp-up and down and 30 s stimulation with the target intensity at 200 μA followed by 14.5 min without stimulation), and EP measures for 120 minutes after stimulation.

**Figure 3.**
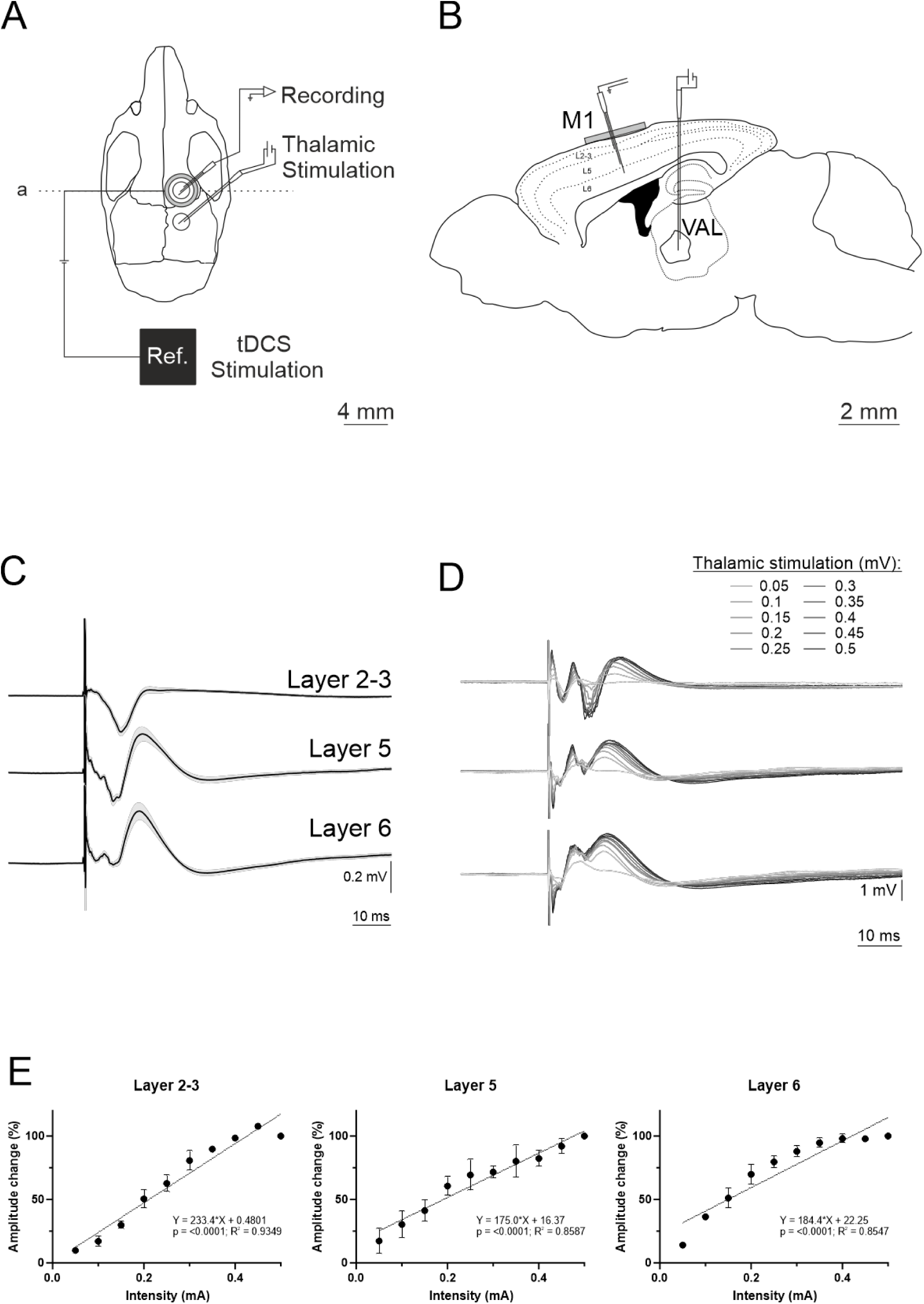
Experimental setup and thalamic EP characterization in M1. A) Experimental setup for concurrent in vivo thalamic stimulation, electrophysiological recordings across layers of M1 and tDCS administration. B) Lateral view showing electrode locations. C) Averaged EP across 10 mice from layer 2-3, layer 5 and layer 6 (ERP, N = 10). D) Superimposed EPs from one mouse recorded at different thalamic stimulation intensities (n = 15 trials for each intensity). E) Quantification of EP amplitude changes at different intensities of thalamic stimulation. Data normalized to the amplitude recorded at 0.5 mA (N = 2 mice). Error bars represent SEM). M1: primary motor cortex; Ref.: tDCS reference electrode.

**Figure 4.**
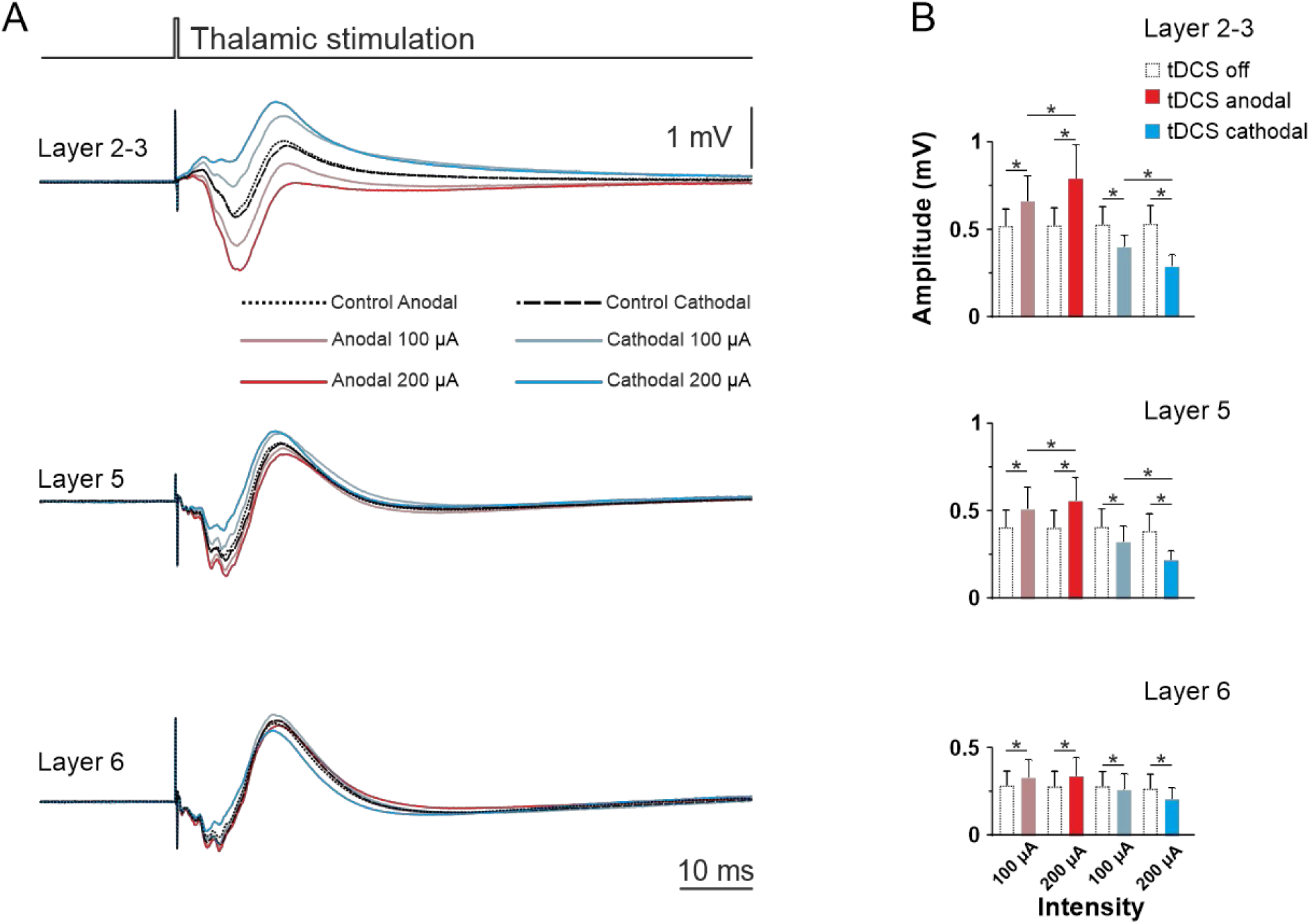
Online effects of 5s tDCS over M1. A) Averaged EPs (n = 15 trials for each intensity) recorded in M1 from a representative animal during the baseline condition before tDCS began (dotted or dashed lines), and superimposed recordings during anodal (red traces) or cathodal (blue traces) tDCS. B) Quantification and statistical analysis of EP amplitudes across the different layers. Empty bars represent values during the baseline period (tDCS OFF) and filled bars represent values during tDCS (tDCS ON) (N = 15 mice; *p < 0.05, LSD). Error bars represent SEM. Asterisks represent statistical differences against baseline (tDCS off).

**Figure 5.**
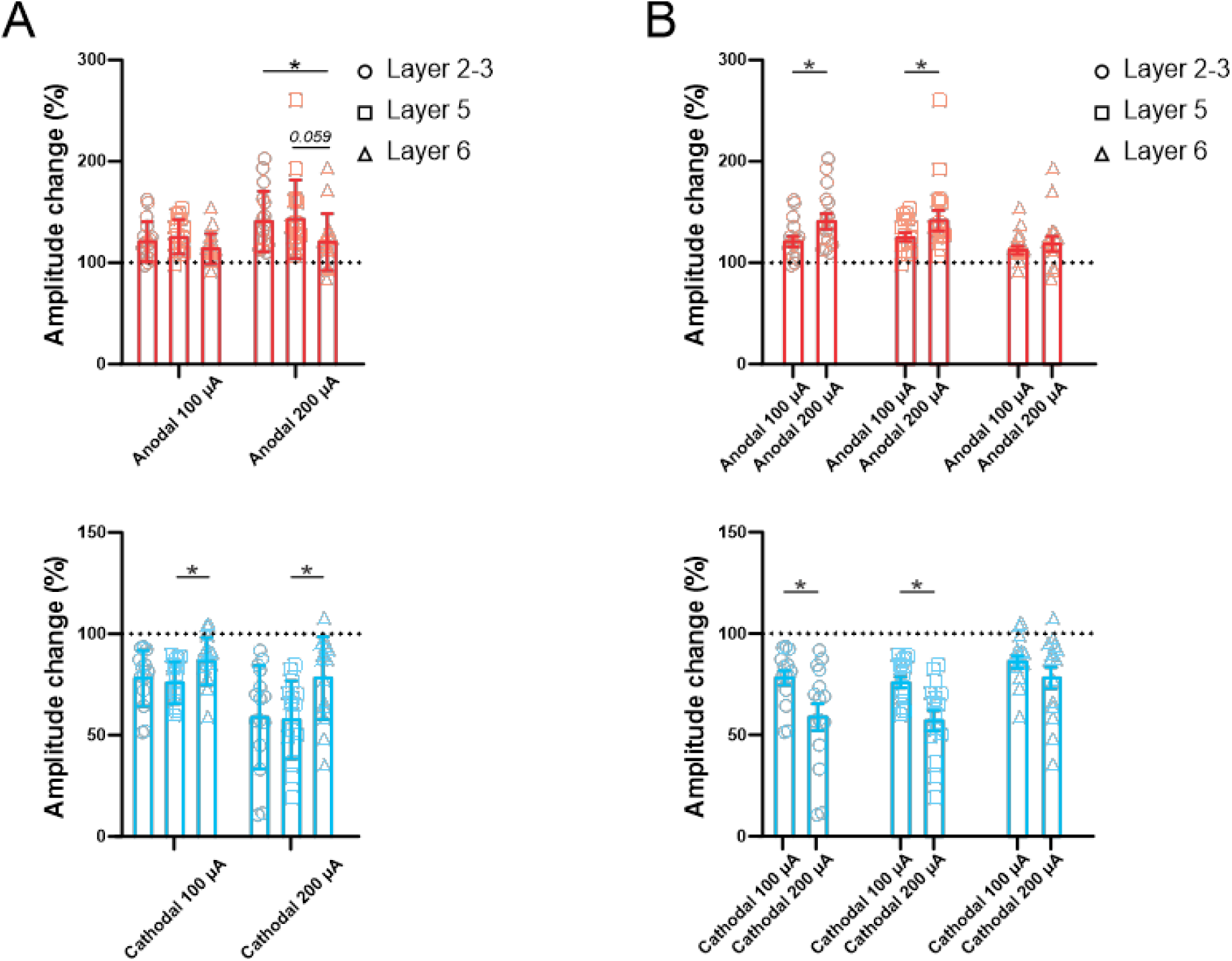
Quantification and statistical analysis of normalized EP amplitudes. A) Normalized amplitude changes (individual data represented by symbols) and statistical comparison between the three different layers for any given tDCS intensity during anodal (top) and cathodal (bottom) stimulation, showing greater modulation in more superficial layers. B) Normalized amplitude change values and statistical comparison between the two different tDCS intensities for any given layer during anodal (top) and cathodal (bottom) stimulation, showing greater modulation for larger intensities. (N = 15 mice). Error bars represent SEM. Asterisks mark statistical differences between layers (A) or between intensities (B), based on post hoc Fisheŕs LSD tests (*p < 0.05).

**Figure 6.**
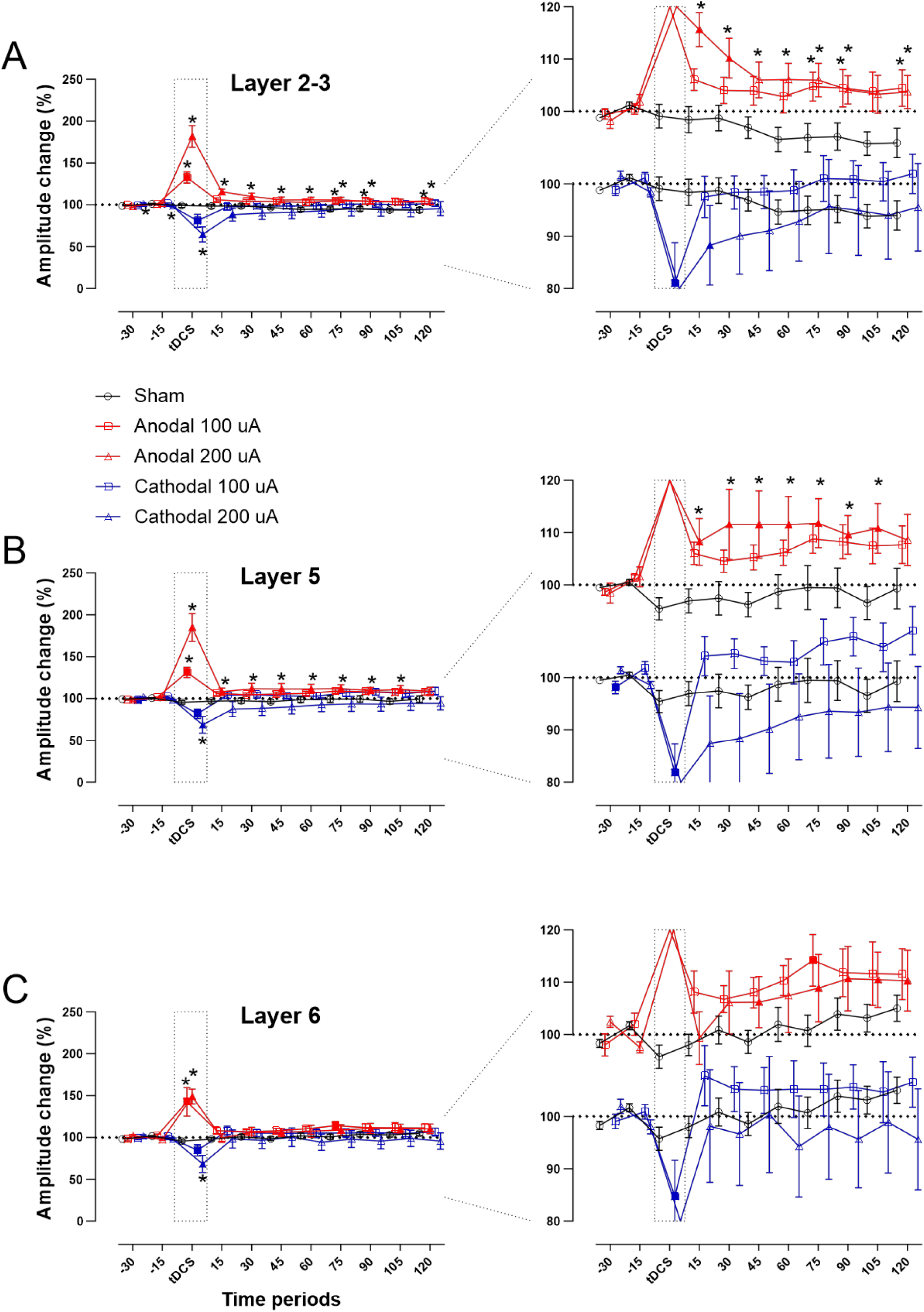
Effects of 15min tDCS over M1 during and after intervention. A,B,C) Normalized EP amplitude changes are depicted for all tDCS conditions for layer 2-3 (A), layer 5 (B) and layer 6 (C) recordings. Red traces for anodal and blue traces for cathodal tDCS with 100 (squares) and 200 (triangles) μA, black traces for sham (circles) tDCS (*p < 0.05, LSD). Error bars represent SEM. Asterisks show statistical differences against the sham condition, filled symbols represent statistical differences against the -15min baseline.10 mice (6 females) were included in the sham condition; N = 11 mice (5 females) in the anodal 100 μA condition; N = 12 mice (6 females) in the anodal 200 μA condition; N = 14 mice (6 females) in the cathodal 100 μA condition; N = 12 mice (4 females) in the cathodal 200 μA condition.

**Table 3.** Overall ANOVA conducted for the after-effects of tDCS. Normalized evoked-potential amplitude values were compared in a 3-factorial mixed-model ANOVA. d.f. = degrees of freedom.

| Factor | d.f. | F-value | p-value | Partial Eta squared |
| --- | --- | --- | --- | --- |
| layer | 1.637* | 1.494 | 0.231 | 0.027 |
| time | 2.754* | 4.576 | 0.005 | 0.078 |
| condition | 4 | 5.001 | 0.002 | 0.270 |
| layer*condition | 6.548* | 0.642 | 0.710 | 0.045 |
| time*condition | 11.017* | 15.532 | <0.001 | 0.535 |
| layer*time | 4.518* | 2.145 | 0.068 | 0.038 |
| layer*time*condition | 18.073* | 1.932 | 0.014 | 0.125 |

Notes: \*: Greenhouse-Geisser corrected dfs

To receive more specific information about the polarity- and intensity-specific effects of stimulation for each layer, we performed a subsequent ANOVA for every polarity separately (Table 4). For anodal stimulation, we observed significant main effects for the factors Time (d.f: 1.934, F = 30.154, p = **<0.001**) and Intensity (d.f: 2, F = 9.633, p = **<0.001**) and the Time*Intensity (d.f: 3.868, F = 11.920, p = **<0.001**), Layer*Time (d.f: 3.325, F = 3.437, p = **0.016**) and Layer*Time*Intensity interactions (d.f: 6.650, F = 2.435, p = **0.026**), whereas for the cathodal polarity only the main effect of Time (d.f: 2.783, F = 9.809, p = **<0.001**) and the Time*Intensity interaction (d.f: 5.566, F = 2.479, p = **0.032**) were significant.

**Table 4.** Secondary ANOVAs conducted for the after-effects of tDCS. Normalized evoked-potential amplitude values were compared in separate 3-factorial mixed-model ANOVAs for each polarity separately. d.f. = degrees of freedom.

| Polarity | Factor | d.f. | F-value | p-value | Partial Eta squared |
| --- | --- | --- | --- | --- | --- |
| Anodal | layer | 1.521* | 0.676 | 0.475 | 0.022 |
|  | time | 1.934* | 30.154 | <0.001 | 0.501 |
|  | intensity | 2 | 9.633 | <0.001 | 0.391 |
|  | layer*intensity | 3.043* | 1.195 | 0.323 | 0.074 |
|  | time*intensity | 3.868* | 11.920 | <0.001 | 0.443 |
|  | layer*time | 3.325* | 3.437 | 0.016 | 0.103 |
|  | layer*time*intensity | 6.650* | 2.435 | 0.026 | 0.140 |
| Cathodal | layer | 1.548* | 1.617 | 0.212 | 0.047 |
|  | time | 2.783* | 9.809 | <0.001 | 0.229 |
|  | intensity | 2 | 1.227 | 0.306 | 0.069 |
|  | layer*intensity | 3.096* | 0.228 | 0.882 | 0.014 |
|  | time*intensity | 5.566* | 2.479 | 0.032 | 0.131 |
|  | layer*time | 3.486* | 0.957 | 0.425 | 0.028 |
|  | layer*time*intensity | 6.972* | 0.831 | 0.563 | 0.048 |

Notes: \*: Greenhouse-Geisser corrected dfs

The post-hoc analysis comparing each 15 min time window of the real vs the sham condition (asterisks in Fig.6), and between each 15 min window and the -15 min time window of baseline (filled symbols in Fig.6) showed two different results during and after stimulation.

During the 15 minutes of stimulation, the EP amplitude was modulated in a similar way as for the online effect experiments (Fig. 6, left), with the highest tDCS intensities increasing and decreasing, for anodal and cathodal tDCS respectively, the amplitude of EP across all layers, and in an intensity-dependent manner (larger EP modulation for larger intensities).

Regarding the after-effects, we observed another pattern of results. Up to 120 minutes after tDCS was switched off (zoom in Fig. 6, right panel), 200 μA anodal tDCS resulted in EP amplitude increases across layers 2-3 (except the penultimate time point) and 5 (except the last time point) in comparison with the sham condition, but in comparison with its own baseline period only layer 5 after 200 μA showed a persistent increase (except the last time point), that for layer 2-3 only lasted for the first 30 minutes. For layer 6, only the comparison with its own baseline showed an amplitude increase after 200 μA anodal tDCS at 45 minutes and during the last hour of the recording session. After 100 μA anodal tDCS, layer 2-3 showed a significant EP amplitude increase in comparison with the sham condition, but only for 3 points at the end of the recording session, and for layer 6 only in comparison with its own baseline at one time point 75 minutes. On the other hand, cathodal stimulation, despite modulating the EP amplitude during stimulation, only resulted in significant aftereffects in the first 15 minutes after 200 μA cathodal tDCS whwere this rgis EP dreased in comparison with its own baseline period in layer 2-3.

These results demonstrate the efficacy of anodal, but only minor effects of cathodal M1-tDCS to induce long lasting changes of neuronal excitability, showing a polarity-, intensity- and layer-dependent modulation for anodal tDCS.

## Discussion

In this study we showed that an *in vivo* awake mouse model of transcranial electrical stimulation is well suited to elicit acute and neuroplastic excitability alterations similar to tDCS effects in humans. This mouse model allows simultaneous electrophysiological recordings and transcranial stimulation and is therefore suitable as a translational approach to test different tDCS protocols like those applied in humans. We observed a polarity-, intensity-, and layer-dependent modulation of neuronal excitability for short duration (5s) transcranial stimulation, with anodal tDCS increasing and cathodal tDCS decreasing neuronal excitability. Moreover, for higher tDCS intensities, a stronger modulation was observed. This modulation was similar during the administration of 15 minutes of tDCS, with anodal tDCS increasing EP amplitudes across all cortical layers in an intensity-dependent manner, and cathodal decreasing EP amplitudes across layers. After tDCS was switched off, we observed that only anodal tDCS at high (200 μA) stimulation intensity produced significant long-lasting after-effects, whereas the after-effects of cathodal tDCS were restricted to 15 min after stimulation in the most superficial layer. In addition, we revealed that the different cortical layers were not modulated uniformly in all cases, with layer 6 being less affected during administration of transcranial currents. Below, we discuss potential mechanisms underlying the observed effects, followed by a discussion of why a mouse model for transcranial stimulation is necessary to understand the impact of these non-invasive brain stimulation techniques on cerebral excitability in larger detail.

The primary motor cortex is one of the brain regions frequently stimulated in non-invasive brain stimulation experiments in humans, partly for the ease to obtain a neurophysiological marker of brain excitability, i.e., MEP. Nonetheless, this is a measurement of the excitability of the corticospinal tract, not a selective, and direct measurement of cortical excitability (Klomjai et al., 2015), and only layer 5 neurons of M1 project directly through the corticospinal tract (Di Lazzaro and Ziemann, 2013), though other elements of the cortical motor circuit are also activated by TMS, and specific TMS protocols are available which are suited to monitor more specifically intracortical inhibition, and facilitation (Wagle-Shukla et al., 2009), including measures of cortical excitability directly via TMS-evoked cortical potentials, which do however not allow to identify layer-specific effects (Mosayebi-Samani et al., 2023). In contrast, animal models allow to selectively and directly asses neuronal excitability, and activity by electrophysiological recordings with high spatial and temporal resolution, for example, demonstrating the physiological effects of transcranial electrical stimulation on spike timing (Krause et al., 2019), LFP oscillations (Krause et al., 2017), the correlation between LFP modulation and learning (Márquez-Ruiz et al., 2012), as well as the involvement of astrocytes or microglial cells in tDCS effects (Monai et al., 2016; Mishima et al., 2019). When we applied tDCS for only 5s (Fig. 4), EP amplitudes increased and decreased for anodal and cathodal tDCS, respectively, in an intensity-dependent manner, i.e., larger stimulation intensities induced more prominent changes, a pattern of modulation also observed in different species (Márquez-Ruiz et al., 2012), including humans (Nitsche and Paulus, 2000; Sugawara et al., 2015; Vaseghi et al., 2015; Agboada et al., 2019; Mosayebi Samani et al., 2019), and in different cerebral regions, including somatosensory (Sánchez-León et al., 2021) and visual (Cambiaghi et al., 2011) cortices. Given the relatively simple cortical geometry of mice, including reduced gyrification, these results are compatible with somatic polarization observed *in vitro* (Bikson et al., 2004; Farahani et al., 2021) and, very recently, also *in vivo (Sánchez-León et al., 2025)*, in a way that anodal stimulation depolarizes the soma of neurons and increases spiking activity while cathodal stimulation hyperpolarizes it, decreasing neuronal firing rate. Another explanation accounting for this phenomenon might be that layer 6 pyramidal neurons are less responsive to polarization due to their size and/or morphology (Radman et al., 2009; Thomson, 2010), and intracortical connectivity in rodents, in which layer 6 pyramidal neurons do not receive excitatory inputs from layer 2/3, in difference to layer 5 neurons. In contrast, pyramidal and inhibitory layer 6 neurons do receive inhibitory input from superficial cortical layers (Briggs, 2010). Further experiments will be needed to answer this question, especially with recent advances in the non-invasive assessment of excitability of deeper cortical layers in humans (Kurz et al., 2019).

Regarding prolonged stimulation protocols, which were expected to induce neuroplastic after-effects, we found a similar modulation during the 15 min of stimulation (Fig. 6, during tDCS) as that observed during 5s stimulation, with anodal tDCS increasing and cathodal tDCS decreasing EP amplitudes in an intensity-dependent manner for anodal stimulation. Nevertheless, when tDCS was switched off, the magnitude of these changes decreased and only after anodal stimulation at 200 µA we observed a significant increase of EP amplitudes that lasted for at least 2 hours after intervention in layers 2-3 and 5, and only after cathodal stimulation at 200 µA for the first 15 minutes in layer 2-3. Explanations for the only minor after-effects after cathodal stimulation could be the variability between individuals, a bottom effect, given that the highest applied tDCS intensity (200 μA cathodal) showed a present, but relatively minor effect on the EP amplitude, the fact that the EP amplitude in the sham condition was not completely stable over time but also decreased gradually, and due to the fact that the mice were not at rest but were allowed to run during the recording, which differs from the respective condition in humans, and might have compromised the hyperpolarizing effect of cathodal tDCS during stimulation. Future studies should explore these candidate factors systematically. Since after intervention, no external currents are applied to the brain, other factors than the externally applied electric field must explain these persistent excitability changes. Here, synaptic modulation through intra-neuronal Ca^2+^ levels (Islam et al., 1995; Nitsche et al., 2003; Mosayebi-Samani et al., 2020), glial cells (Monai et al., 2016), the brain-derived neurotrophic factor (BDNF) (Fritsch et al., 2010; Ranieri et al., 2012), different glutamate receptors such as NMDA (Nitsche et al., 2003, 2004), mGluR5 (Sun et al., 2016), AMPA (Stafford et al., 2018; Martins et al., 2019), and adenosine (Márquez-Ruiz et al., 2012) has been proposed.

One of the advantages of this animal model for transcranial stimulation is the direct assessment of M1 excitability, probably the most frequently targeted brain region in human tDCS experiments. Intracortical recordings allowed us to discover a layer-dependent modulation of M1, a phenomenon also observed *in vivo* in the somatosensory cortex of mice and *in vitro* in mice and the human M1 (Sun et al., 2020), suggesting that tDCS could have a different impact on different layers in other regions of the cortex. Our tDCS model also has the advantage of allowing awake animal experiments in combination with electrophysiological recordings. Usually, *in vivo* animal studies exploring tDCS effects on neuronal activity have been performed under anesthesia (Lynn Bindman et al., 1964; Cambiaghi et al., 2011; Vöröslakos et al., 2018; Kunori and Takashima, 2019; Asan et al., 2020; Gellner et al., 2020), but different anesthetic drugs are probably affecting tDCS-induced plasticity via their influence on neuronal excitability, activity, and membrane polarization, as shown for the voltage-dependent sodium channel blocker carbamazepine, which blocks acute and after-effects of anodal tDCS in humans (Nitsche et al., 2003), and might thus be confounding factors in these experiments.

On the other hand, animal models have some limitations. It has been shown in humans that TMS effects on M1 excitability depend on the behavioral state (active vs resting), something hard to control in awake mice that could partially explain the absence of after-effects of cathodal stimulation (Kahl et al., 2022). In addition, we are aware that in our model we applied higher current densities (1.05 and 2.11 mA/cm^2^ for 100 and 200 μA) than those typically applied in humans (0.028 mA/cm^2^ for 1 mA and an electrode size of 35 cm^2^).

Therefore, it is important to take these differences into account when considering a generalization to human studies. Another important limitation of the current study is the measurement of local field potentials, which reflects the synchronized activity of many neurons nearby the recording electrode. Future experiments should take advantage of single neuron recordings with electrophysiological methods for high temporal resolution and with calcium imaging to resolve the spatial distribution of the neurons being modulated as well as their identity, i.e., the different types of excitatory or inhibitory neurons.

Our findings show that a mouse model for M1 non-invasive neuromodulation can replicate some of the effects observed in human experiments, positioning it as a promising candidate to bridge the gap between *in vitro* animal studies and human experiments, by allowing a direct characterization of neuronal excitability with better spatial and temporal resolution in animals, and translating this knowledge to human tDCS studies. It also opens the door for future experiments to explore the mechanisms behind tDCS effects to a larger degree, as could be the exploration of effects at a single neuron level, the direct measurement of intracellular Ca^2+^ levels, or the characterization of the molecular pathways causing the after-effects.

## Acknowledgements

**MAN reveived support from the** German Centre for Mental Health (DZPG), Bochum 44789, Germany, and from the German Research Foundation Research Unit 5429/1 (467143400), NI 683/17-1

## Author contributions

C.A.S-L, C.V.T. and M.A.N conceived the original idea and designed the experiments. C.A.S-L performed the experiments. C.A.S-L and P.A.R-V performed the data analysis. C.A.S-L, C.V.T. and M.A.N wrote the paper. All authors contributed to the final edition of the manuscript.

## Declaration of interests

The authors declare the following financial interests/personal relationships which may be considered as potential competing interests: Michael A. Nitsche is a member of the scientific advisory boards of Neuroelectrics and Precisis. None of the remaining authors have potential conflicts of interest to disclose.

## Notes

### Competing Interest Statement

The authors have declared no competing interest.

